# Single-molecule analysis of gap and nick binding by LIG1 and LIG3α at the final step of DNA repair

**DOI:** 10.1101/2025.06.25.661597

**Authors:** Surajit Chatterjee, Kar Men Lee, Kanal Balu, Melike Çağlayan

## Abstract

DNA ligase (LIG) 1 and LIG3α repair broken single-strand breaks in the phosphodiester backbone at the final ligation step of DNA excision repair pathways, and complement each other during nuclear replication in case of unligated Okazaki fragments. We previously reported that both ligases discriminate against nicks containing non-canonical ends and ligate gap intermediate if left unfilled by DNA polymerases. However, it remains unknown how the dynamics of DNA binding differ for gap *versus* nick substrates by LIG1 and LIG3α at single-molecule level. Here, using total internal reflection fluorescence (TIRF) and ligation assays, we showed that LIG3α binds less frequently but forms longer-lived complex than LIG1 for nicks containing canonical A:T, mismatch G:T, and damaged 8oxoG:A, and they exhibit subtle differences in discriminating unusual ends. Moreover, our results identified gap DNA as a new target to which LIG1 and LIG3α can bind as efficient as their preferential nick sites. We showed gap ligation and observed that more percentage of LIG1 molecules form stable long-lived complex on DNA containing one nucleotide gap, whereas LIG3α forms short-lived gap complex without any differences in the percentage of molecule forming gap-bound complex. Finally, our findings demonstrated that LIG1 can still stably bind to larger gaps with better recognition, whereas LIG3α binding becomes further infrequent and shorter-lived. Overall, our study provides single-molecule insights into intricate differences between LIG1 and LIG3α for binding to a range of mutagenic and deleterious DNA repair and replication intermediates that could be a threat for maintaining genome stability at the final step.

## Introduction

Genomic DNA is susceptible to damage by a variety of endogenous sources and environmental exposures such as radiation, toxins, air pollution, and sunlight (1). As a consequence, tens of thousands of DNA strand breaks are made continually and naturally throughout the genome (2). Single-strand breaks (SSBs) are the most common form of DNA damage arising at least once per cell every 1 or 2 sec (3). Oxidative stress has been implicated in the formation of SSBs directly via the disintegration of the oxidised base/nucleotide by reactive oxygen or nitrogen species (4). Other common sources of SSBs include unligated Okazaki fragments of DNA replication and the intermediates of DNA excision repair pathways such as base excision repair (BER), nucleotide excision repair (NER), and mismatch repair (MMR) (5). Unrepaired SSBs can lead to persistent breaks and tend to be converted into toxic and lethal double-strand breaks (3–5). DNA ligases join broken SSBs in the phosphodiester backbone during critical DNA transactions (6). The conversion of nicks into a phosphodiester bond by DNA ligase requires a rapid recognition and binding to avoid deterioration of the exposed DNA ends to exonuclease degradation and to eliminate the possibility for an increased frequency of recombination (7). Yet, the mechanism driving nick search and binding by human DNA ligase remain a critical knowledge gap in our understanding the molecular determinants of faithful ligation during DNA replication and repair.

The three human ligase genes (*LIG1*, *LIG3* and *LIG4*) encode ATP-dependent DNA ligases (8). DNA ligase 1 (LIG1) is the main replicative ligase playing a critical role for maturation of Okazaki fragments with >50 million ligation events during DNA replication, acts in a fusion of sister chromatids by targeting double-stranded DNA breaks, and is responsible for the majority of DNA ligase activity in proliferating cells (9–13). The reiteration of mutations identified in the *LIG1* gene of human individuals (P529L, R641L, R771W), whose symptoms included developmental abnormalities, immunodeficiency and lymphoma, in a mouse model has proven that the LIG1 deficiency causes genetic instability and predisposition to cancer (14–16). DNA ligase 3α (LIG3α) is the vital ligase for mitochondrial genome integrity and its deficiency underlies abnormal mitochondrial function as reported in Alzheimer’s disease and related dementias (17,18). Furthermore, LIG3α plays a critical role for SSB repair in coordination with X-ray repair cross-complementing protein 1 (XRCC1) and Poly [ADP-ribose] polymerase 1 (PARP1) (19,20). LIG1 and LIG3α finalize an ultimate ligation step at the end of DNA excision repair pathways (21). Therefore, elucidating the mechanism by which LIG1 and LIG3α recognize their targets and efficiently bind to DNA to ensure faithful joining of broken strand breaks is critical for understanding how genome stability is maintained during DNA repair and nuclear replication.

LIG1 and LIG3α share a highly conserved catalytic core consisting of the adenylation (AdD) and the oligomer binding (OBD) domains (22). While the catalytic activity of the enzyme is largely governed by the active site residues (K568 of LIG1 and K421 of LIG3α) residing in the AdD domain, the N-terminal α-helical extension next to the catalytic core is recognized as DNA binding domain (DBD) that binds to a nick more robustly (23). In addition to highly conserved catalytic core, LIG1 and LIG3α contain flanking and unrelated amino (N)- and/or carboxyl (C)-terminal regions that mediate their interactions with other repair and replication proteins, and direct them for their participation in different DNA transactions in nucleus or mitochondria (24–28). In our recently reported single-molecule study (29), we demonstrated that LIG1 is enriched near nick sites and the protein exhibits diffusive behavior to form a long-lived ligase-nick complex after binding to a non-nick region, yet LIG1 C-terminal mutant (lacking non-catalytic N-terminal region) predominantly binds throughout DNA non-specifically to the regions lacking a nick site for shorter time.

LIG1 and LIG3α catalyze the formation of a phosphodiester bond by in-line nucleophilic attack of 3’-hydroxyl (OH) on 5’-phosphate (PO_4_) to seal adjacent DNA ends in a highly conserved ligation reaction (22). Yet, they fail in case of modified ends at the nick (30). Our previous studies demonstrated that LIG1 and LIG3α fail in the presence of mismatched or damaged base at the 3’-end of nick after incorporation of non-canonical mismatches or oxidized nucleotides (*i.e*., 8-oxodGTP) by DNA polymerase β (polβ) at the downstream steps of BER pathway (31–38). This results in abortive ligation and the formation of 5’-adenylated (AMP) intermediate, which is itself very serious damage at the nick and could become a persistent DNA strand break if left until being processed by DNA end processing enzymes such as Aprataxin (39). We further reported at atomic resolution in X-ray crystallography studies of LIG1 that the ligase active site employs distinct strategies while engaging with nicks containing mismatched or oxidatively damaged ends (40–43). For example, LIG1 favors G:T or deters A:C mismatch for end joining and engages with nick harboring an oxidative damage depending on the dual coding potential of 8oxoG(*anti*):C(*anti*) and 8oxoG(*syn*):A(*anti*) that forms non-mutagenic Watson-Crick and mutagenic Hoogsteen base pairing, respectively. However, how the presence of canonical *versus* damage or mismatch-containing ends at the nick site could affect initial DNA binding modes of LIG1 and LIG3α at single-molecule level remains entirely unknown.

In our previous studies (42,43), we also demonstrated that the presence of ribonucleotides confounds the downstream steps of BER pathway where gaps left unfilled by polβ can be subsequently ligated by LIG1 and LIG3α. Furthermore, we reported that a defective gap filling by polβ cancer-associated variants (*i.e.,* gastric carcinoma E295K) that were identified in a high percentage of tumors and exhibit aberrant nucleotide incorporation activity even in the presence of canonical nucleotides. Similarly, unfilled gaps by these polβ variants results in subsequent ligation of gap intermediate by LIG1 and LIG3α at the final step of BER pathway (44,45). This ligation of one nucleotide gap is polβ and free nucleotide independent so the gap ligation could be also a consequence even in the presence of other repair polymerases. Yet, the binding dynamics of LIG1 and LIG3α to these deleterious repair intermediates containing gap sites over regular nick targets remains unknown.

Recent studies revealed that unfilled ssDNA gaps could be formed in the lagging strand in case of unligated Okazaki fragments if not rapidly processed by the canonical DNA replication machinery due to pertubrated LIG1 and Flap Endonuclease 1 (FEN1) functions, which leads to PARP1 activation (46,47). It has been also demonstrated that PARP1’s auto-PARylation recruits LIG3α and polβ to process and fill gaps during the mechanism that complements the canonical replication pathway where LIG3α can back-up LIG1 deficiency, which is fundamental to the completion of lagging strand synthesis in unperturbed cells and requires the BRCA-RAD51 pathway (48). Furthermore, these structures could be formed during NER where an initial excision of a damage leaves a gap in the DNA template that is then filled in by DNA polymerase and ligated to the downstream phosphodiester backbone by LIG1 or LIG3α to complete the repair process (49,50). Defects in gap filling (*i.e*., enlargement of the excision gap as reported in human cells following exposure to UV radiation or other related agents) have the potential to be mutagenic and lethal (51). Yet, DNA binding characteristics of LIG1 and LIG3α to these larger gap structures has never studied at single-molecule level.

In the present study, by employing a total internal reflection fluorescence (TIRF) microscopy combined with ligation assays *in vitro*, we extensively characterized DNA binding dynamics of LIG1 *versus* LIG3α and compared the lifetimes of bound and unbound states for a range of nick substrates containing ligatable and canonical A:T, mismatched G:T, or damaged 8oxoG:A ends in real-time. Our findings demonstrated that LIG3α forms longer-lived complex and exhibits longer unbound time than LIG1 for all nicks. Furthermore, both ligases show short-lived binding to DNA in the absence of nick site. In the ligation assays, we obtained subtle differences in nick sealing efficiencies of LIG1 and LIG3α for all 12 possible non-canonical mismatches depending on the architecture of 3’-primer:template terminus and mutagenic ligation of 3’-8oxodG:A, yet LIG3α exhibits relatively slower end joining and lower amount of ligation products for all nicks. We also investigated DNA binding modes of LIG1 and LIG3α to dsDNA containing gap site(s). For one nucleotide gap, compared to a single nick site, more LIG1 molecules form longer-lived complex characteristristic of target recognition, whereas the binding lifetime of LIG3α decreases without any changes in percentage of long-bound molecules. Furthermore, our findings with larger gap (five nucleotides) demonstrated similar LIG1 binding as nick and shorter gap, whereas LIG3α binding becomes further infrequent and shorter-lived compared to nick and one nucleotide gap. Overall, our study provides a mechanistic inshight into DNA binding mechanism at the ligation step and reveales important differences between LIG1 and LIG3α during their interaction with nick and gap substrates that could be formed when DNA polymerases incorporate mismatches or damaged nucleotides and left gaps unfilled leading to the formation of a range of deleterious and lethal repair and replication intermediates, which could adversely impact the maintenance of genome integrity.

## Methods

### Protein purifications

LIG3α (pET-24b) protein was purified as described (29–44). Briefly, the protein was overexpressed in BL21(DE3) *E. coli* cells in Terrific Broth (TB) media with kanamycin (50 μgml^−1^) and chloramphenicol (34 μgml^−1^) at 37 °C. Once the OD_600_ reached 1.0, the cells were induced with 0.5 mM isopropyl β-D-thiogalactoside (IPTG) and overexpression continued for overnight at 20 °C. The cells were then grown overnight at 16 °C. After centrifugation, the cells were lysed in the lysis buffer containing 50 mM Tris-HCl (pH 7.0), 500 mM NaCl, 20 mM imidazole, 2 mM β-mercaptoethanol, 5% glycerol, and 1 mM phenylmethylsulfonyl fluoride (PMSF) by sonication at 4 °C. The lysate was pelleted at 16,000 x rpm for 1 h and then clarified by centrifugation and filtration. The supernatant was loaded on a HisTrap HP column and purified with an increasing imidazole gradient (0-300 mM) elution at 4 °C. The collected fractions including his-tag LIG3α protein were then subsequently loaded on a HiTrap Heparin column and eluted with a linear gradient of NaCl up to 1 M. The protein was finally purified by a Superdex 200 Increase 10/300 column in the buffer containing 50 mM Tris-HCl (pH 7.0), 500 mM NaCl, 5% glycerol, and 1 mM DTT. LIG3α active site mutant K421A was purified similarly with the full-length wild-type protein (Supplementary Scheme 1). LIG1 (pET-24b) protein was purified as described (29–44). Briefly, the protein was overexpressed in BL21(DE3) *E. coli* cells in TB media at 37 °C for 8 h and induced with IPTG. The cells were harvested, lysed at 4 °C, and then clarified as described above. The supernatant was loaded on a HisTrap HP column and purified with an increasing imidazole gradient (0-300 mM) elution at 4 °C. The collected fractions were then subsequently loaded on a HiTrap Heparin column with a linear gradient of NaCl up to 1 M. His-tag LIG1 protein was then further purified by a Superdex 200 Increase 10/300 column in the buffer containing 50 mM Tris-HCl (pH 7.0), 500 mM NaCl, 5% glycerol, and 1 mM DTT. LIG1 EE/AA (E346A/E592A) and active site K568A mutants were purified similarly with the full-length wild-type protein (Supplementary Scheme 1). DNA polymerase (pol) β (pET-28a) protein was purified as described (29–44). Briefly, the protein was overexpressed in BL21(DE3) *E. coli* cells in TB media at 37 °C and induced with IPTG. The cells were then grown overnight at 16 °C. The cells were harvested, lysed at 4 °C, and then clarified as described above. The supernatant was loaded on a HisTrap HP column and purified with an increasing imidazole gradient (0-300 mM) elution at 4 °C. The collected fractions including polβ protein were then subsequently loaded on a HiTrap Heparin column and eluted with a linear gradient of NaCl up to 1 M. All proteins used in this study were dialyzed against storage buffer containing 25 mM TrisHCl (pH 7.4), 100 mM KCl, 1 mM TCEP, and 10% glycerol, concentrated, frozen in liquid nitrogen, and stored at -80 °C in aliquots.

### Single-molecule gap and nick DNA binding measurements

Using a total internal reflection fluorescence (TIRF) microscope (Nikon Eclipse Ti2-E), we performed single-molecule imaging experiments to visualize gap and nick DNA bindings by LIG1 and LIG3α as previously reported (29). For this purpose, we labeled both ligases with Cy5 after their pre-adenylation and used dsDNA substrate (34-mer) containing 5’-Biotin and 3’-AF488 label (29). These dsDNA substrates contain no nick site, a single nick site harboring ligatable Watson-Crick base paired A:T, mismatched G:T, or damaged 8oxoG:A at the 3’-end as well as gap with one- or five-nucleotides (Supplementary Tables 1 and 2).

Briefly, glass coverslips were functionalized with a mixture of biotin-PEG-SVA and mPEG-SVA and microfluidic channels were assembled using the passivated coverslips and rinsed three times with the T50 buffer containing 10 mM Tris-HCl (pH 8.0) and 50 mM NaCl. Streptavidin (0.2 mg/ml) in the T50 buffer was then flowed onto the slide, reacted with the biotin-PEG for 5 min, and then washed with the T50 buffer. AF488 labeled DNA substrate was diluted to a final concentration of 10 pM in the imaging buffer containing 1 mM HEPES (pH 7.4), 20 mM NaCl, 0.02% BSA (w/v), and 0.002% Tween 20 (v/v), flowed onto the slide, and allowed to incubate for 3 minutes for immobilization. Excess unbound DNA substrate was then washed by flowing with the imaging buffer. To reduce non-specific surface binding of the labeled proteins, we further passivated the slide surface by incubating with 10 mg/ml BSA for 10 minutes. Oxygen-scavenging system (OSS) consisting of 44 mM glucose, 165 U/ml glucose oxidase from Aspergillus niger, catalase (2,170 U/ml) as well as 10 mM Trolox were added to slow photobleaching and to reduce photo blinking, respectively. Finally, LIG1^Cy5^ and LIG3α^Cy5^ proteins (1 nM) in the imaging buffer along with the OSS system were flowed onto the slide and allowed to equilibrate before imaging with an objective-based TIRF microscope. Both AF488 and Cy5 dyes were simultaneously excited using ∼5 mW (at the source) of 488 nm and 640 nm lasers, respectively. Emissions from two fluorophores were separated into two channels using a Cairn Optosplit III image splitter and simultaneously recorded at 100 ms time resolution using a Hamamatsu SCMOS camera using NIS-Elements acquisition software (Nikon, version: AR 6.02.01).

For data analysis, the locations of molecules and fluorophore intensity over time traces were extracted from the raw movie files using Nikon NIS-Elements analysis software (Nikon, version: AR 6.02.01). Genuine fluorescence time traces for individual molecules were selected as described previously (29). Selected time traces were idealized using a two-state hidden Markov model (HMM) for the unbound and bound states in QuB and the rastergrams summarizing several individual traces were generated from the individual trace HMMs using custom written MATLAB script. Cumulative frequency of the bound and unbound dwell-time distributions was plotted and fitted in Origin Lab (version 2024b) with single or double exponential functions to obtain the bound (t_bound_) and unbound (t_unbound_) states lifetimes.

### DNA ligation assays

Nick DNA substrates with a 6-carboxyfluorescein (FAM) label were used in the ligation assays to test nick sealing efficiency of LIG1 and LIG3α (Supplementary Scheme 2). Nick DNA substrates containing Watson-Crick base paired ends, and all possible 12 non-canonical mismatches were prepared by annealing upstream oligonucleotides 3’-dA, 3’-dT, 3’-dG, or 3’-dC with template oligonucleotides A, T, G, or C (Supplementary Table 3). Nick DNA substrates containing oxidative DNA damage were prepared by annealing upstream oligonucleotide 3’-8oxodG with template oligonucleotides A or C (Supplementary Table 3). DNA ligation assays were performed as described (29–44). Briefly, the reaction mixture contains 50 mM Tris-HCl (pH 7.5), 100 mM KCl, 10 mM MgCl_2_, 1 mM ATP, 1 mM DTT, 100 µg ml^-1^ BSA, 1% glycerol, and DNA substrate (500 nM) in the final volume of 10 µl. The reaction was initiated by the addition of LIG1 or LIG3α (100 nM) and incubated at 37 °C for the time points as indicated in the figure legends. The reactions were stopped by mixing with an equal volume of loading dye containing 95% formamide, 20 mM ethylenediaminetetraacetic acid, 0.02% bromophenol blue, and 0.02% xylene cyanol. Reaction products were separated by electrophoresis on an 18% Urea-PAGE gel, the gels were scanned with Typhoon PhosphorImager RGB, and the data was analyzed using ImageQuant software. The ligation assays were performed similarly for mutants (LIG1 K568A, LIG1 EE/AA, LIG3α K421A) with the wild-type full-length proteins.

### Gap ligation assays

Gap ligation assays were performed as previously reported (29–44). One nucleotide gap DNA substrate with template base C was used (Supplementary Table 4). The reaction mixture contains 50 mM Tris-HCl (pH 7.5), 100 mM KCl, 10 mM MgCl_2_, 1 mM ATP, 1 mM DTT, 100 µg ml^-1^ BSA, 1% glycerol, and DNA substrate (500 nM) in the final volume of 10 µl. The reaction was initiated by the addition of LIG1 or LIG3α, and the reaction mixture was incubated at 37 °C for the time points as indicated in the figure legends. The reaction products were then mixed with an equal amount of gel loading buffer containing 95% formamide, 20 mM EDTA, 0.02% bromophenol blue, and 0.02% xylene cyanol and separated by electrophoresis on 18% Urea-PAGE gel. The gels were finally scanned and the data were analyzed as described above. In the control reaction, we tested a complete ligation of resulting nick product after polβ dGTP insertion opposite C in gap DNA in the reaction mixture containing 50 mM Tris-HCl (pH 7.5), 100 mM KCl, 10 mM MgCl_2_, 1 mM ATP, 1 mM DTT, 100 µg ml^-1^ BSA, 1% glycerol, dGTP (100 µM), and DNA substrate (500 nM) in the final volume of 10 µl. In this case, the reaction was initiated by the addition of a pre-incubated enzyme mixture of polβ/LIG3α and the reaction mixture was incubated at 37 °C for the time points as indicated in the figure legends. The gels were finally scanned and the data were analyzed as described above.

## Results

### Nick DNA binding by LIG1 and LIG3α at single-molecule level

We employed single-molecule fluorescence co-localization approach using TIRF microscopy to visualize and compare nick DNA binding characteristics of LIG1 and LIG3α in real-time. The 5’-biotinylated and AF488-labeled dsDNA substrate contains a single nick site and harbors ligatable canonical (A:T), mismatch (G:T), or damage (8oxoG:A) at the 3’-end. These nick DNA substrates were immobilized on a biotinylated glass slide utilizing the multivalency of streptavidin and the real-time LIG1 and LIG3α binding to nick DNA were monitored (Figure 1A). For nick with canonical A:T end, the analyses of individual fluorescence time trajectories show repeated transient Cy5 co-localizations with AF488 within a diffraction-limited spot, indicating dynamic binding of LIG1 and LIG3α on the DNA (Figure 1B-C). Rastergrams generated from several individual traces idealized by hidden Markov model demonstated similar short and long binding events for both ligases (Figure 1D-E). In the presence of nicks containing G:T and 8oxoG:A mismatches, the individual fluorescence time trajectories and rastergrams also show charisteristic nick DNA binding for both LIG1 and LIG3α (Supplementary Figures 1-2). However, the binding events are noticeably less frequent for LIG3α than that of LIG1. Our results demonstrated differences in binding life-times for all nicks containing canonical *versus* non-canonical ends (Supplementary Figure 3). In our previous study (29), we reported that the long binding events (∼8 sec) are due to the formation of stable LIG1-DNA complex at nick site, whereas the short binding events are due to the non-specific binding of the protein to non-nick regions. In the present study, we further demonstrated that LIG1 can bind to nick containing canonical or mismatched ends with similar efficiency where t_bound_ and t_unbound_ times do not show significant difference among all nicks (Figure 2A,C). However, for LIG3α, when compared to LIG1, the bound times are moderately higher for nicks containing 8oxoG:A (1.5 times) than G:T mismatch (1.3 times) ends, whereas the unbound time is significantly higher (1.7 times). These results suggest that the initial interactions of LIG1 and LIG3α with nick DNA are similar in terms of overall binding parameters although there are differences in the stability of nick-bound complexes and their ability to recognize a nick site.

**Figure 1.**
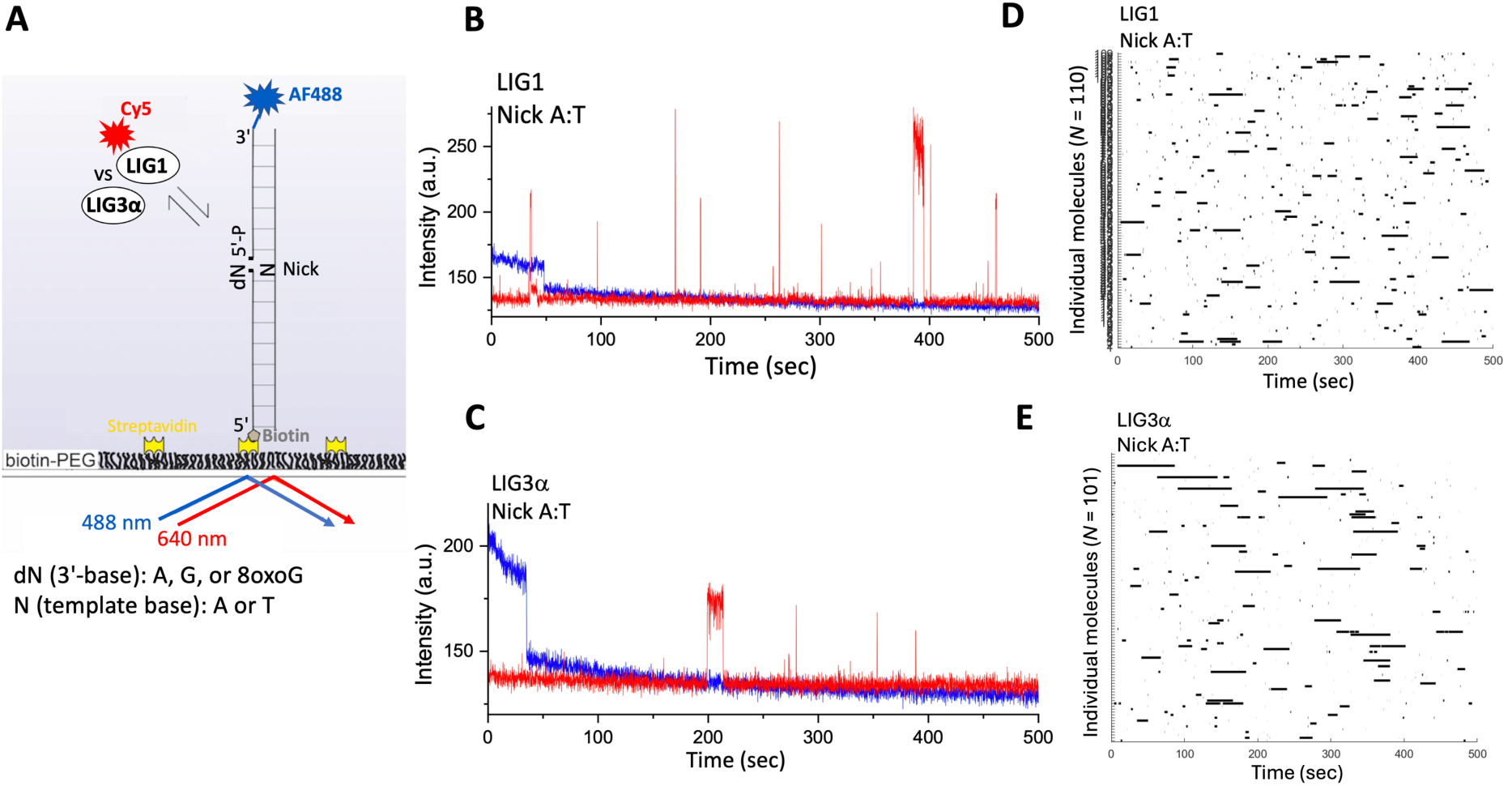
Single-molecule characterization of nick DNA binding by LIG1 and LIG3α. **(A)** Scheme shows 3’-AF488-labeled dsDNA immobilized on PEG-coated and biotinylated slide surface to monitor Cy5-labeled LIG1 or LIG3α binding to nick DNA containing A:T, G:T, and 8oxoG:A at 3’-end. **(B-C)** Fluorescence intensity *versus* time traces show repeated protein binding events of LIG1 and LIG3α to dsDNA containing a nick with A:T end. (**D-E**) Rastergrams for individual molecules from selected traces are shown for LIG1 and LIG3α proteins displaying the nick DNA binding behavior.

**Figure 2.**
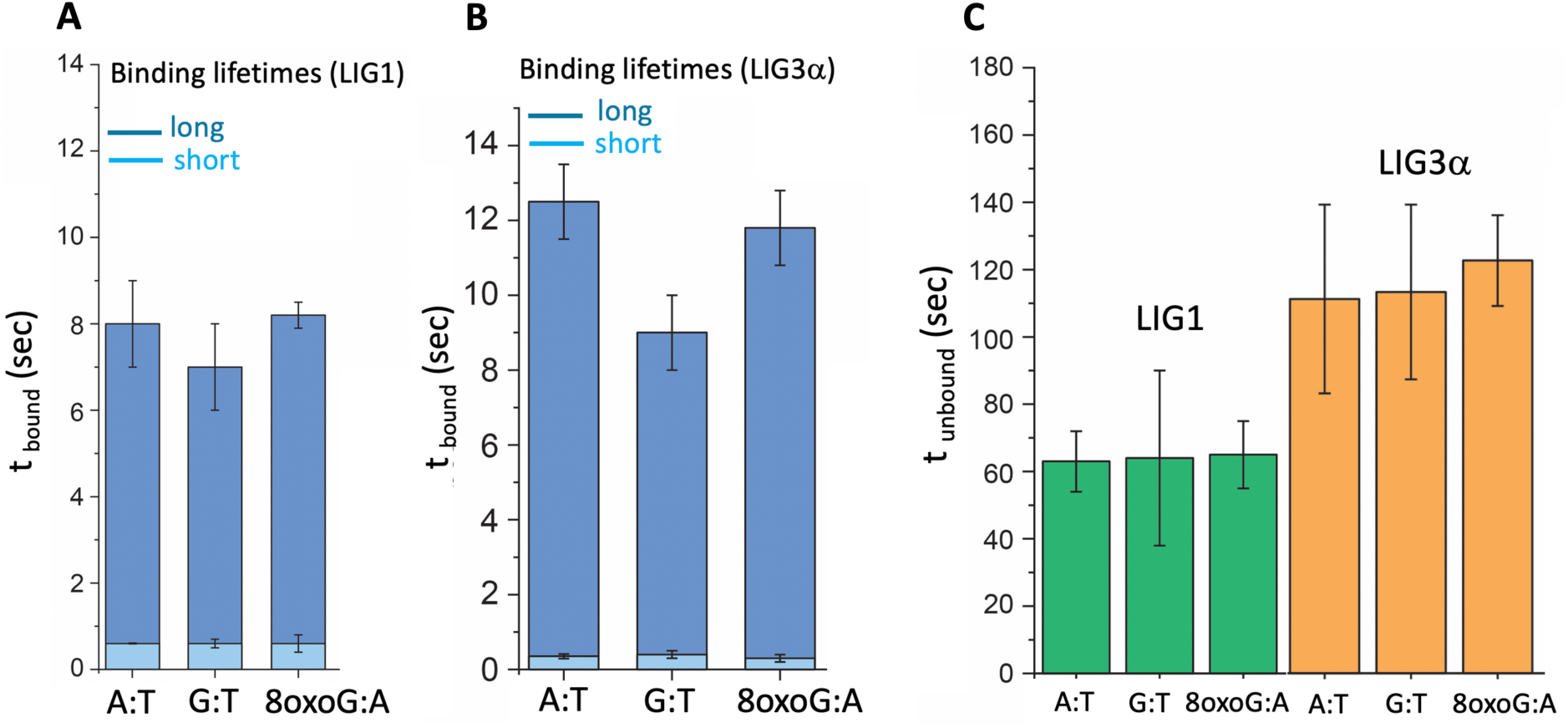
Single-molecule DNA binding characterization of LIG1 and LIG3α for nicks containing canonical or mismatched ends. **(A-C)** Bar graphs show the differences in the protein-bound (*t*_bound_) state (A-B) and protein-unbound (*t_un_*_bound_) state (C) that represent the binding lifetimes of LIG1 and LIG3α to nick containing A:T, G:T, and 8oxoG:A at the 3’-end.

We also performed single-molecule assays in the presence of dsDNA without a nick site that mimics an undamaged DNA or a final ligation product after sealing of both DNA ends by a DNA ligase (Figure 3). Our results showed similar binding behaviours of LIG1 and LIG3α as indicated by the individual fluorescence time trajectories and demonstrated that both ligases exhibit very short-lived and rare binding behaviours (Supplementary Figures 4-5). We then investigated LIG1 K568A and LIG3α K421A active site mutants to compare their binding life-times with corresponding wild-type proteins (Figure 4). Our results showed significant decrease in bound-times for both mutants, whereas LIG3α K421A shows ∼4 times difference compared to LIG1 K568A (∼2 times). Moreover, the unbound-time for LIG1 K568A remains same as wild-type, whereas it is significantly increased from 107 ± 27 sec (wild-type) to 174 ± 6 sec for LIG3α K421A mutant (Supplementary Figures 6-7). Overall, these results suggest that both active site mutants cannot form a stable nick-ligase complex during initial DNA binding and the mutation in LIG3α that diminihes pre-adenylation of the enzyme has more prominent in minimizing the ability of the ligase to find a nick site.

**Figure 3.**
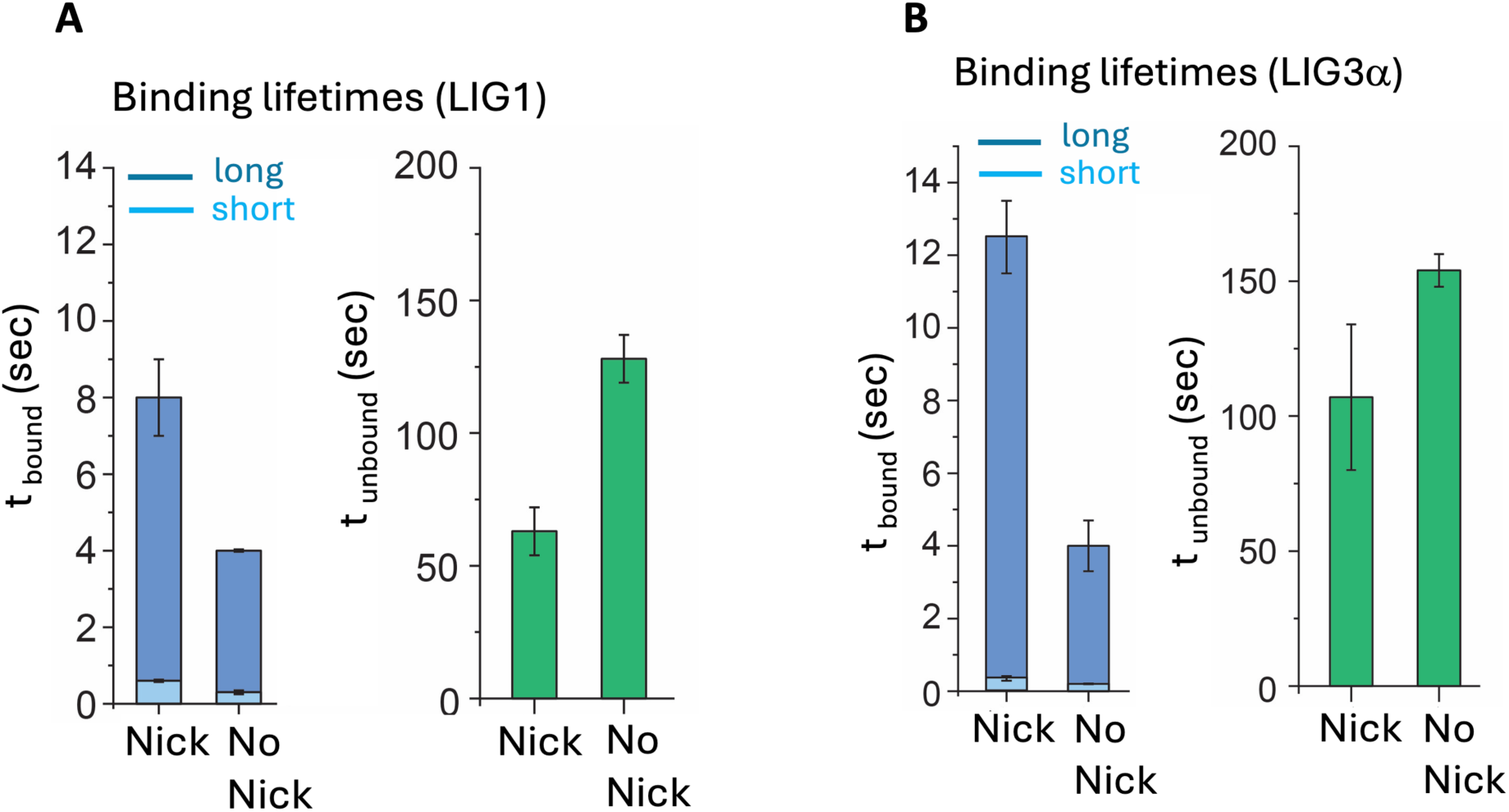
Single-molecule DNA binding characterization of LIG1 and LIG3α in the absence and presence of nick site. **(A-B)** Bar graphs show the difference in the protein-bound (*t*_bound_) and unbound (*t_un_*_bound_) states that represent the binding lifetimes of LIG1 and LIG3α to dsDNA with and withour a single nick site.

**Figure 4.**
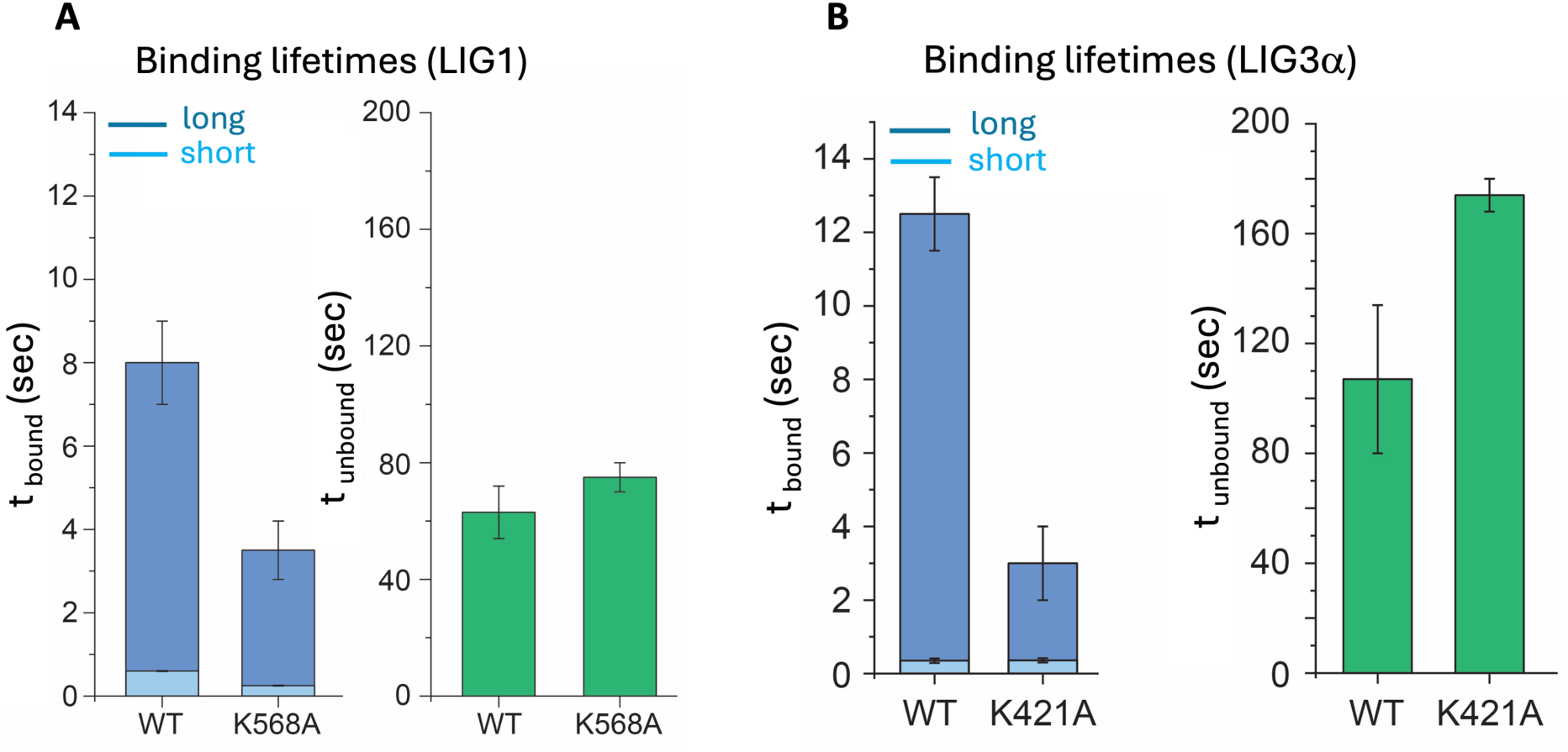
Comparison of nick DNA binding life-times by LIG1 and LIG3α active site mutants. **(A-B)** Bar graphs show the difference in the protein-bound (*t*_bound_) and unbound (*t_un_*_bound_) states that represent the nick DNA binding lifetimes of LIG1 K568A and LIG3α K421A active site mutants.

The fidelity of LIG1 is mediated by the active site residues (E346 and E592) and the mutations at those side chains (E346A/E592A or EE/AA) result in the formation of a cavity at the ligase active site leading to low-fidelity enzyme (40–43). We further investigated nick binding mode of LIG1 using the EE/AA mutant for substrates containing canonical (A:T) and damaged (8oxoG:A) ends. Interestingly, in both cases, we observed longer bound-time compared to the wild-type protein, suggesting the formation of more stable nick complex by LIG1 EE/AA low-fidelity mutant (Figure 5). The unbound-times were found to be similar for both nick substrates and for both LIG1 proteins, suggesting that the mutation that impacts the fidelity of the ligase has no significant effect on the ability of the protein to find a nick site (Supplementary Figures 8-9).

**Figure 5.**
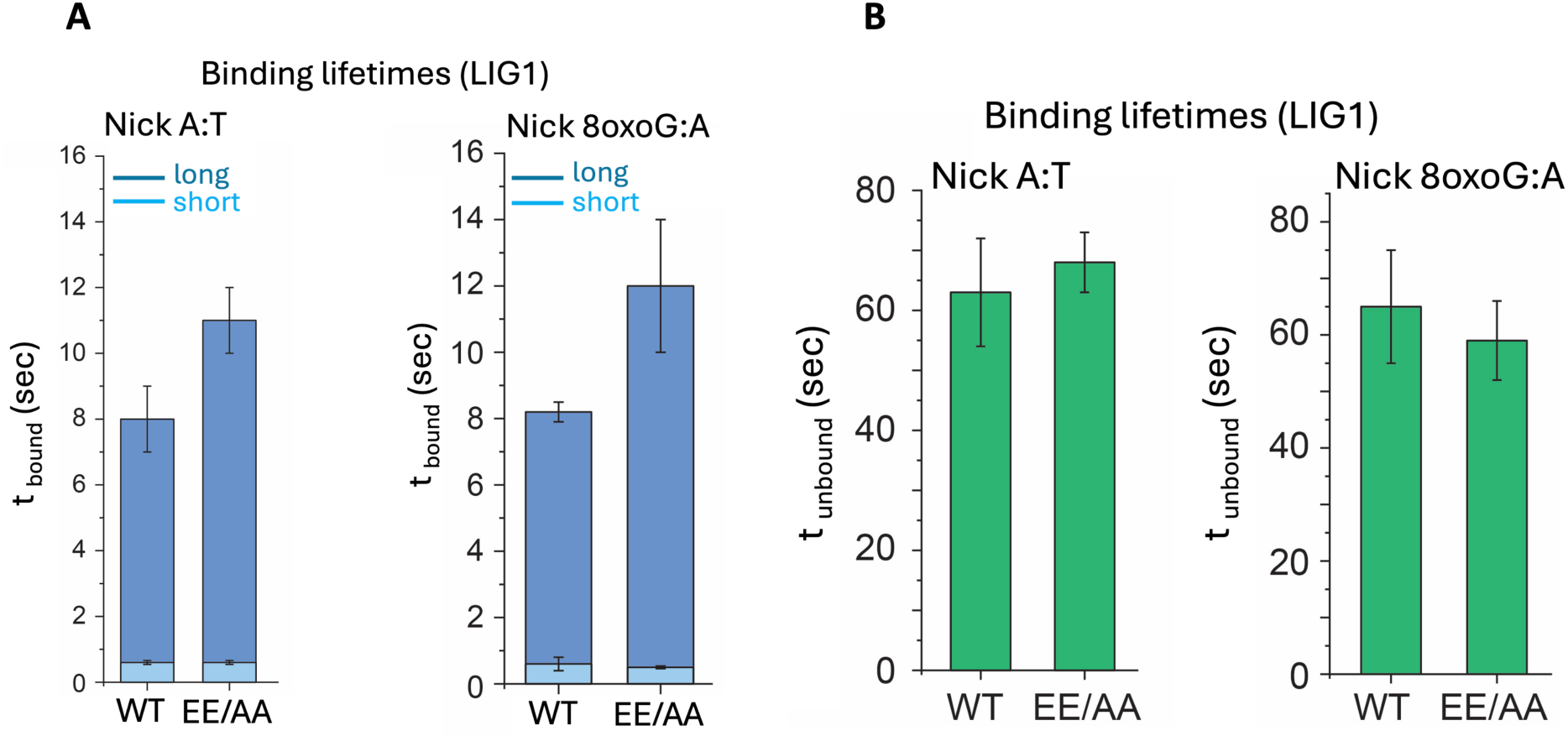
Comparison of nick DNA binding life-times by LIG1 wild-type and low-fidelity mutant. **(A-B)** Bar graphs show the difference in the protein-bound (*t*_bound_) and unbound (*t_un_*_bound_) states that represent the nick DNA binding lifetimes of LIG1 wild-type and EE/AA low-fidelity mutant.

### Substrate specificity of LIG1 and LIG3α for nicks containing canonical, mismatched, or damaged ends

In addition to nick DNA bindings at single-molecule level, we comprehensively characterized substrate specificity of LIG1 and LIG3α in the ligation assays *in vitro* for nick DNA substrates containing all possible 12 non-canonical mismatches and 8oxoG at the 3’-end.

The ligation efficiency of nicks containing mismatch-containig ends demonstrated that LIG1 and LIG3α exhibit subtle differences depending on the architecture of 3’-terminus:template base pairing (Supplementary Figures 10-13). For template A mismatches, LIG1 shows efficient nick sealing of 3’-dA:A and 3’-dC:A mismatches, while LIG3α discriminates against all mismatches, and no ligation product was observed in the presence of 3’-dG:A by both ligases (Supplementary Figure 10). LIG1 indiscriminately ligates all possible template T mismatches, 3’-dT:T, 3’-dC:T, 3’-dG:T, and we obtained ligation products by LIG3α in the presence of 3’-dC:T and 3’-dG:T mismatches, which were relatively less efficient (Supplementary Figure 11). In the presence of template G mismatches, we obtained no ligation product for 3’-dA:G and 3’-dG:A mismatches by LIG1 and LIG3α (Supplementary Figure 12). LIG1 was found relatively more efficient for sealing of template C mismatches and we observed a time-dependent increase in the amount of ligation products in the presence of 3’-dT:C mismatch by both ligases (Supplementary Figure 13). In the presence of the nick substrate containing damaged ends, our results showed the mutagenic ligation of 3’-8oxoG:A by LIG1 and LIG3α and the end joining ability of both ligases was significantly diminished when 8oxoG is paired with template C (Supplementary Figure 14). For all four canonical nick DNA substrates containing Watson-Crick base paired ends, we obtained an efficient ligation by LIG1 and a reduced nick sealing by LIG3α particularly at earlier time points of ligation reaction (Supplementary Figures 15-16).

As expected, there was no DNA ligation by LIG1 K568A and LIG3α K421A active site mutants (Supplementary Figure 17). LIG1 low-fidelity mutant EE/AA shows relatively more efficient nick sealing for all substrates containing 3’-mismatches compared to the wild-type enzyme (Supplementary Figures 18-19). In the presence of nick DNA substrate containing 3’-8oxoG, we observed less ligation failure products by the effect of E346A and E592A mutations (Supplementary Figure 20), while the ligation of canonical ends was as efficient as LIG1 wild-type (Supplementary Figure 21).

### Gap DNA binding by LIG1 and LIG3α at single-molecule level

In addition to nick DNA bindings with canonical *versus* mismatched or damaged ends, we employed single-molecule fluorescence co-localization approach to visualize gap DNA binding by LIG1 and LIG3α. We first investigated the dynamics of both ligases on AF488-labeled dsDNA harboring one nucleotide gap that was immobilized on a biotinylated glass slide utilizing the multivalency of streptavidin (Figure 6A). This substrate mimics unfilled gap intermediate by DNA polymerase during DNA synthesis step of DNA excision repair pathway as we previously reported (44).

**Figure 6.**
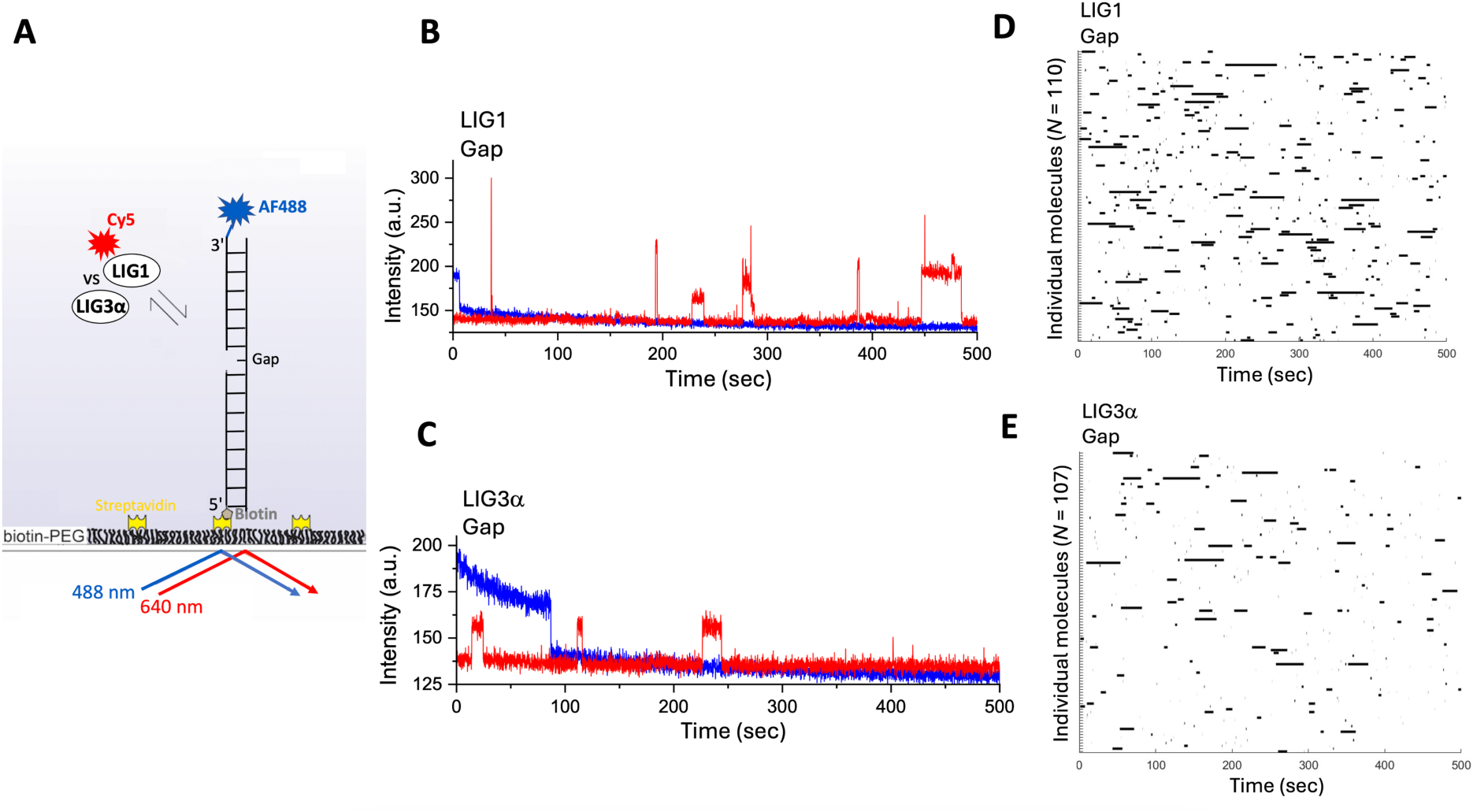
Single-molecule characterization of gap DNA binding by LIG1 and LIG3α. **(A)** Scheme shows 3’-AF488-labeled dsDNA immobilized on the PEG-coated and biotinylated slide surface to monitor Cy5-labeled LIG1 or LIG3α binding to one nucleotide gap DNA. **(B-C)** Fluorescence intensity *versus* time traces show repeated protein binding events of LIG1 and LIG3α to gap site. (**C-D**) Rastergrams for individual molecules from the selected traces are shown for LIG1 and LIG3α proteins displaying gap DNA binding behavior.

Our results showed similar gap binding characteristics of LIG1 and LIG3α as shown by the co-localization of AF488 and Cy5 signals within a diffraction-limited spot (Figure 6B-E). Regarding the comparision of nick *versus* gap DNA binding modes, we showed that LIG1 exhibits similar bound and unbound lifetimes (Supplementary Figure 22). However, the percentage of LIG1 molecules depicting longer-lived complex is higher (∼60%) for binding to gap than nick (∼30%), suggesting that more number of LIG1 molecules can recognize gap site on dsDNA and form a stable bound complex (Figure 7A). In contrary, for LIG3α, the lifetime of the longer-lived complex is decreased from 12.5 ± 1 sec (nick) to 8.1 ± 0.4 sec (gap), while the percentage of LIG3α molecules is similar between nick and gap substrates, suggesting that LIG3α has less ability to form stable gap-bound complex on dsDNA (Figure 7B). Furthermore, we measured gap DNA binding in the presence of Mg^2+^, and observed extremely short-lived and less frequent binding for both LIG1 and LIG3α (Supplementary Figures 23-24). Interestingly, these results were found similar as observed for undamaged DNA without a nick (or gap) site (Figure 8). This similarity in terms of short binding by LIG1 and LIG3α to gap DNA in the presence of Mg^2+^ and to DNA in the absence of nick suggests that both ligases can seal one nucleotide gap.

**Figure 7.**
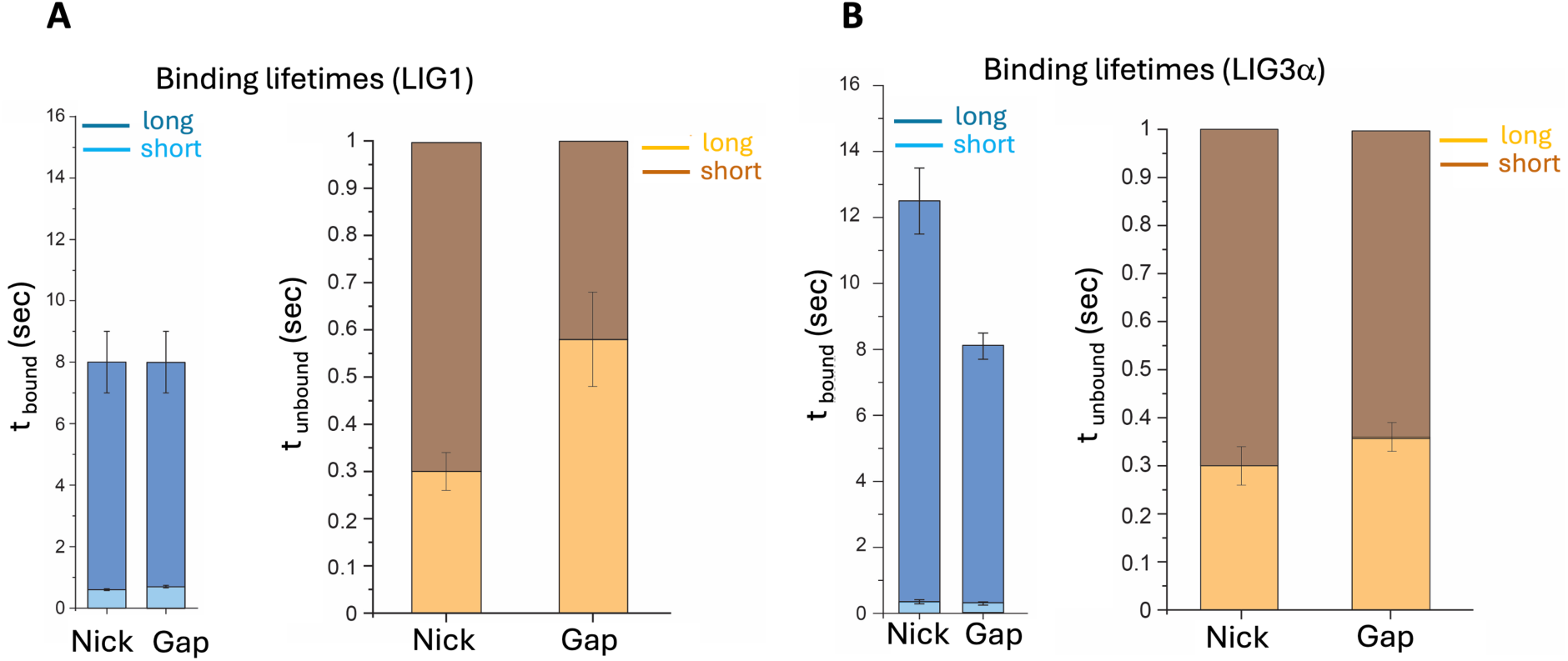
Comparison of nick *versus* gap DNA binding life-times by LIG1 and LIG3α. **(A-B)** Bar graphs show the difference in the protein-bound (*t*_bound_) and unbound (*t_un_*_bound_) states that represent the binding lifetimes of LIG1 and LIG3α to dsDNA containing a single nick site or one nucleotide gap.

**Figure 8.**
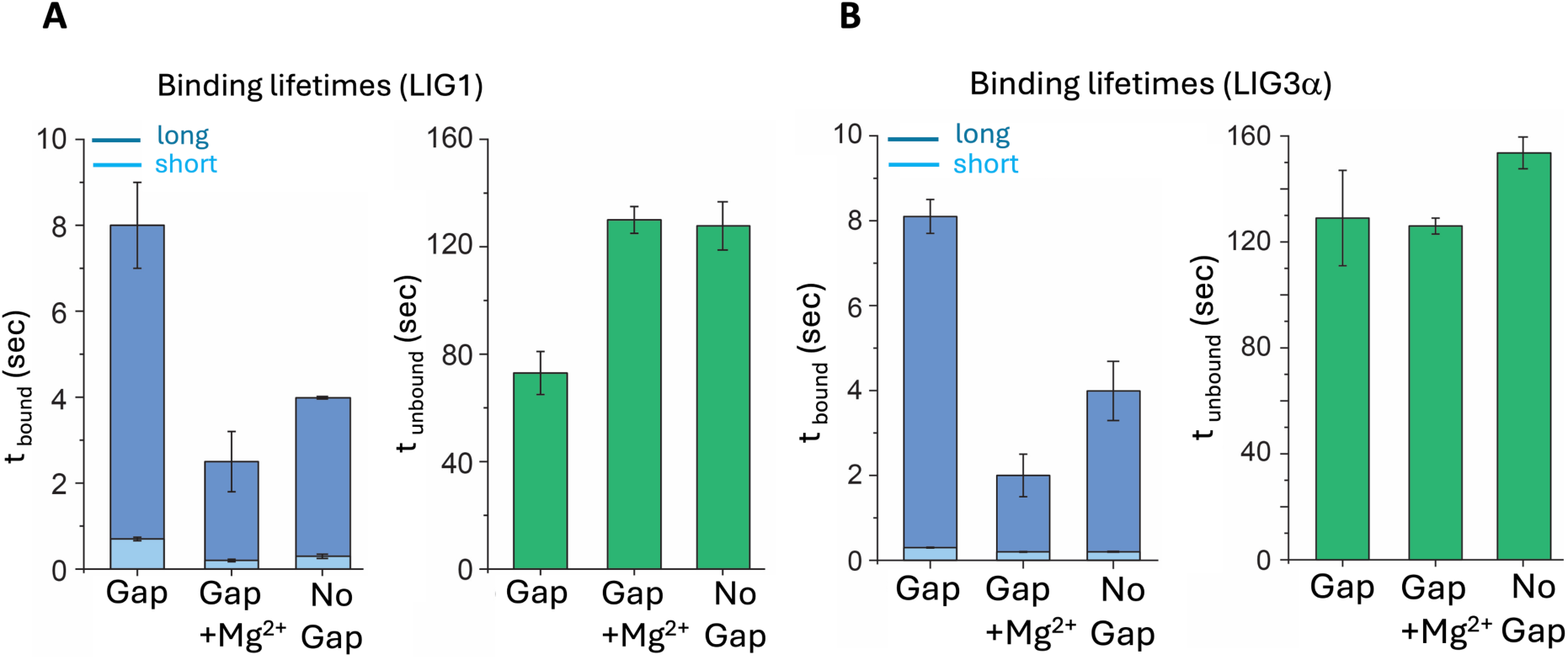
Comparison of gap DNA binding life-times by LIG1 and LIG3α in the presence of a metal ion. **(A-B)** Bar graphs show the difference in the protein-bound (*t*_bound_) and unbound (*t_un_*_bound_) states that represent the binding lifetimes of LIG1 and LIG3α to dsDNA without a nick or gap site *versus* dsDNA containing a gap site in the absence and presence of Mg^2+^.

In the ligation assays, we also demonstrated that LIG1 and LIG3α can ligate one nucleotide gap DNA itself, referred to as gap ligation, in the reactions including DNA ligase and one nucleotide gap DNA substrate (Supplementary Figure 25A). We showed gap *versus* nick ligation as revealed by the difference in the size of the products in the control reactions (Supplementary Figure 25B, line 2 for a complete ligation *versus* lanes 3-8 for gap ligation by LIG1 and lanes 9-14 for gap ligation by LIG3α). In addition, we showed the products of polβ dGTP insertion opposite C in one nucleotide gap DNA substrate as well as the conversion of these dGTP:C insertion products to a complete ligation products over reaction time in the coupled assay including polβ, LIG3α, dGTP, and gap DNA substrate (Supplementary Figure 26).

Finally, we measured DNA binding dynamics of LIG1 and LIG3α in the presence of larger gap (Figure 9). These substrate mimics the replication intermediates that can be formed when SSBs left unfilled on the lagging strand during nuclear replication or the intermediates of NER pathway when post-incision gap filling is interrupted during the repair of UV photoproducts, highlighting such gap structures as potential sources of genome instability (46–51).

**Figure 9.**
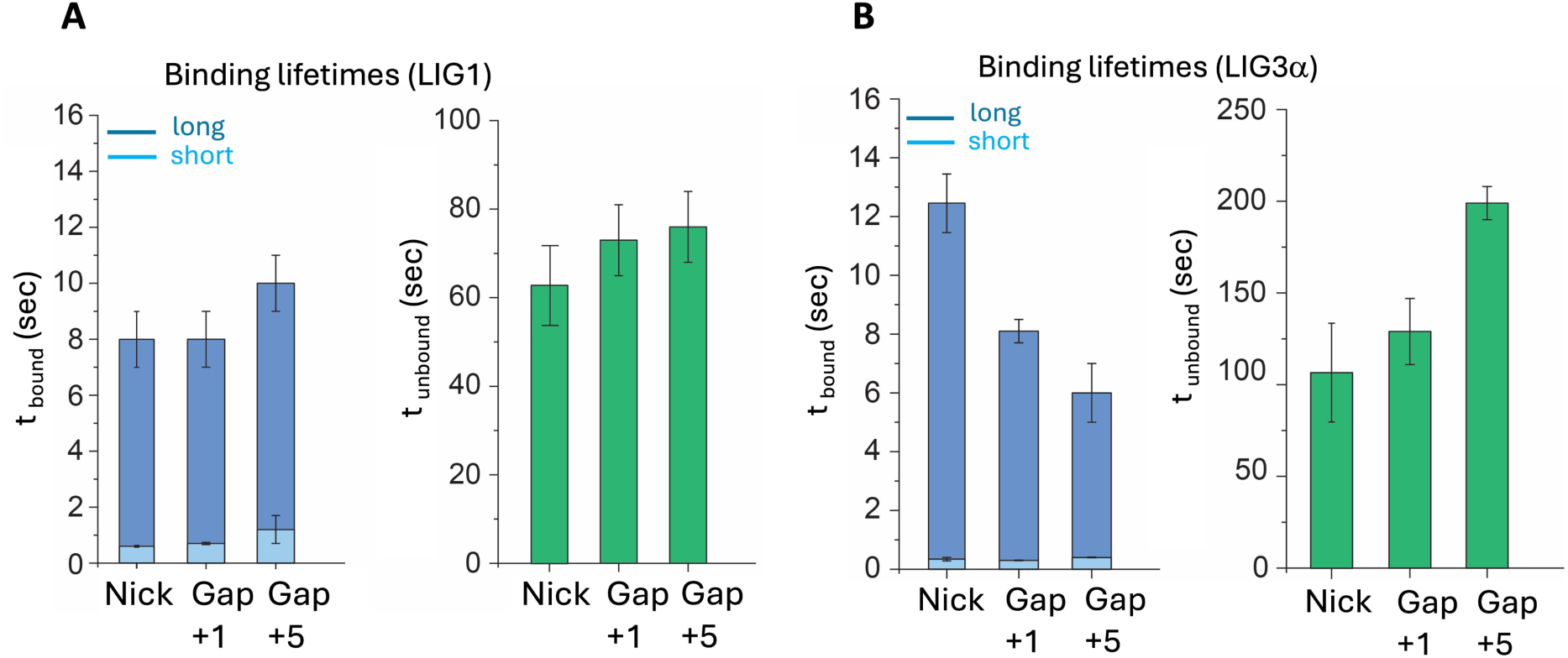
Comparison of binding life-times for nick *versus* short and large gap DNA bindings by LIG1 and LIG3α. **(A-B)** Bar graphs show the difference in the protein-bound (*t*_bound_) and unbound (*t_un_*_bound_) states for LIG1 (A) and LIG3α (B) binding to dsDNA substrates containing a single nick, one- and five- nucleotide gap sites.

Interestingly, our results showed an efficient gap binding by LIG1, with similar bound, unbound times and percentages as observed for nick and one nucleotide gap substrates. Whereas, for LIG3α, further decrease in bound time was observed (6 ± 1 sec) along with significant increase in unbound time (199 ± 9 sec), suggesting less affinity of LIG3α to bind to larger gaps (Supplementary Figures 27 and 28). Together, this data highlights the role LIG1 to bind gap substrates with high affinity, while LIG3α binding is more specific to nick sites.

## Discussion

Mammalian genomes can be exposed to multiple types of DNA damage that can be generated during endogenous cellular processes or by the effect of exogenous sources such as environmental factors including UV-light, chemical toxins, irradiation, inflammation, and nutritional factors (1,2). SSBs can occur as natural intermediates during DNA replication, cytosine demethylation, differentiation, gene rearrangement in the immune system and germ cell development (3). Failure to seal nascent DNA strands results in the formation of potentially deleterious nick in the DNA backbone, making the position vulnerable to exonuclease degradation, increased frequency of recombination, and formation of double-strand breaks (4). Otherwise, these strand breaks can cause loss or addition of genetic information at the break site (7). The joining of SSBs by DNA ligase is essential for maintaining overall genome stability as they have overlapping functions in nucleus and mitochondria (6,8,22). LIG1 and LIG3α join broken SSBs using the complementary strand of the double helix as a template for the formation of a phosphodiester bond between 3’-OH and 5’-PO_4_ ends of nick to create an uninterrepted DNA strand during DNA replication and repair (30,33). It’s also important to note that both ligases can complement each other during nuclear replication and DNA repair (52). Yet, how they efficiently bind to DNA in order to repair broken strand breaks during such criticial DNA transactions to maintain genome integrity is still not fully understood, particularly at single-molecule level.

It is well established that genomic instability drives the progression of a normal cell into a cancer cell and the maintenance of genomic integrity requires high fidelity in all DNA transactions including DNA replication, recombination, and repair (5). LIG1 and LIG3α discriminate a mismatched or damaged base at the 3’-end of nick (22). However, it’s stil unknown whether nicks containing Watson-Crick base-paired *versus* non-canonical ends impact the efficiency of DNA binding.

In our recent single molecule work, we showed that LIG1 binding to undamaged DNA is rare and very short-lived, while in the presence of a nick, we observed long binding events that are characteristics of nick-ligase complex. The binding mechanism of many DNA repair proteins are consists of multiple pathways such as direct binding of the protein from solution to the target site, binding on an undamaged region and followed by sliding or hopping to the target site (53–55). For example, thymine DNA glycosylase (TDG), in addition to its preferred G:T substrate, can switch between binding modes to search different base modifications and have different binding lifetimes dependent on the sequence and downstream repair process (56). We previously showed that LIG1 can bind to a nick site through both non-specific binding to a non-nick site followed by diffusion to nick site, and specific binding to the nick site to form a stable nick-bound complex (29). The bound and unbound times measured from our single molecue data also provide multiple information about the binding mechanism where the short (<2 sec) and long (∼8 sec) bindings suggest the non-target (non-specific) and target site (specific) binding, respectively. On the other hand, the unbound time suggests how fast or slow the ligase finds a target site. As an another important parameter, the percentage of long-bound protein indicates the number of protein molecules that remain stably bound at a target site. In our study, we systematically compared these parameters to elucidate the intricate differences in gap *versus* nick binding characteristics between LIG1 and LIG3α. Our single-molecule measurements demonstrated similar bound and unbound life-times for nick repair intermediates containing A:T, G:T or 8oxoG:A ends that mimic DNA polymerase-mediated incorporation products of mismatch or damaged nucleotides, suggesting that the configuration of 3’-terminus:template base pairing does not impact initial nick DNA binding by LIG1 and LIG3α. However, the binding events were noticeably less frequent for LIG3α, suggesting less affinity to target DNA than that of LIG1. Consistently, our ligation assays demonstrated differences in nick sealing efficiency of all possible 12 non-canonical mismatches and 8oxoG by LIG3α. For example, we obtained ∼4 to 8-fold reduced amount of ligation products for nicks with G:T mismatch, while there was ∼80-fold decrease in mutagenic nick sealing efficiency of 3’-8oxoG:A by LIG3α, which could be explained due to relatively higher unbound time compared to LIG1. We also observed relatively more efficient ligation of mismatches by LIG1 EE/AA mutant compared to the wild-type enzyme, which is consistent with the single-molecule DNA bindings showing longer bound-time to all nicks by the low-fidelity mutant, suggesting the formation of more stable nick complex leading to an increase in ability to join DNA ends.

We further demonstrated, for the first-time at single-molecule level, that LIG1 and LIG3α can efficiently bind to DNA containing a gap site and there are distinct differences in their binding characteristics. For LIG1, the bound and unbound lifetimes are similar for all substrates (nick and short or larger gaps), but more ligase molecule remain bound to gap site compared to nick. This suggests that either more LIG1 molecules can differentiate the gap, compared to nick, from the non-damaged sites on the DNA or, after binding, the protein undergoes certain conformational change that stabilizes the gap-bound complex. In contrary to LIG1, for LIG3α, the lifetime of longer-lived complex gradually decreased from nick to a single to larger gap DNA substrates while the unbound time gradually increases, suggesting that LIG3α has less affinity towards gap sites on DNA and form less stable gap-bound complex than nick. Furthermore, in the presence of Mg^2+^, both ligases show rare and short-lived binding events to gap as observed for intact dsDNA that mimics a sealed nick repair product, demonstrating that LIG1 and LIG3α can bind and ligate one nucleotide gap substrate facilitated by the metal ion. Our previous studies showed that both LIG1 and LIG3α can bind to one nucleotide gap with similar binding affinity as polα and can consequently results in gap ligation (34,44). Single molecule particle tracking measurements in *E. coli* also demonstrated that the polymerase and ligase molecules has less affinity for undamaged DNA however become transiently bind to repair randomly distributed gap and nick sites and that the polymerase has higher nick-translation activity and compete with DNA ligase for nick sealing (57). Given the fact that other types of DNA excision repair pathways such as as ribonucleotide excision repair and mismatch repair also induce SSBs indirectly, gap DNA binding ability of both ligases could generate deleterious repair intermediates and nucleotide deletion products that could be more detrimental than an embedded one due to potential disruptions in the DNA sequence. Overall, these gap structures that can be recognized as target to bind by LIG1 and LIG3α during DNA replication and repair could be a threat for maintaining genome integrity.

Studies have demonstrated that a perturbed gap filling synthesis in NER causes histone H2AX phosphorylation in human quiescent cells and the resultant ssDNA-gap intermediates initiate the phosphorylation of histone H2AX in an ATR-dependent manner (49–51). Similarly, ssDNA gaps remain in the lagging strand when canonical Okazaki fragment processing is incomplete where LIG3α can complement a deficiency in the main replicative ligase, LIG1, for maturation of replication intermediates (46–48). Our findings from single-molecule measurements with DNA substrates containing gap sites demonstrated that although both LIG1 and LIG3α can bind to a single and larger gaps, LIG1 has better ability to recognize and form stable bound complex at the gap sites. This suggest that LIG1 might be the first responder to these gap structures and its deficiency in case of severe immunodeficiency-associated mutations in the *LIG1* gene leading to perturbation in its activity can be complemented by LIG3α.

Overall, we comprehensively characterized the dynamics of nick DNA binding by LIG1 and LIG3α at single-molecule level, which could contribute to understand their binding modes to a range of repair and replication intermediates with unusual ends. We also identified a new target and specific DNA binding properties of both ligases with their upstream gap intermediates that are indeed regular substrates of DNA polymerases. It has been reported that both ligases operate through their interactions with repair and replication protein partners (24–28). For example, through N-terminal non-catalytic and unstructured region, LIG1 interacts with PCNA, replication factor A (RFA), and Rad9-Hus1-Rad1 (9-1-1) clamp complex as well as BER proteins polβ and AP-Endonuclease 1 (APE1) (24–26,34). Similarly, LIG3α forms a repair complex with XRCC1 via C-terminal BRCT domains and N-terminal extension containing a zinc-finger (ZnF) domain plays role in nick DNA binding as a “nick sensing” that increases its sensitivity to SSBs and mediates PARP1 interaction (27,28). Further single-molecule studies are needed to elucidate the mechanism by which intermolecular interactions regulate target search and binding by LIG1 and LIG3α.

DNA ligase inhibitors were identified *in silico* screen based on DNA binding pocket of LIG1 following its atomic resolution structure in complex with nick (58–60). Recent studies have reported an interplay between PARP1/2, PAR synthesis and DNA ligases in genome maintenance (61). Furthermore, PARP and LIG1 or LIG3α inhibitors has been widely used in combined theraphy for leukemia and cancer cells that are defective in homology-dependent repair pathway (62–64). Our findings from single-molecule DNA binding characteristics of LIG1 and LIG3α could contribute to the development and utilization of more effective DNA ligase inhibitors particularly by targeting gap and nick binding abilities of LIG1 or LIG3α in the absence and presence of a damage or mismatch during lagging strand replication and PARP1/2-mediated SSB repair in cancer cells.

## Data Availability

Information and requests of materials used in this research should be directed to Melike Çaglayan.

